# Masculinity and the mechanisms of human self-domestication

**DOI:** 10.1101/143875

**Authors:** Ben Thomas Gleeson

## Abstract

**Objectives:** Pre-historic decline in human craniofacial masculinity has been proposed as evidence of selection for elevated sociability and a process of ‘human self-domestication’ thought to have promoted complex capacities including language, culture, and cumulative technological development. This follows experimental observation of similar changes in non-human animals under selection for reduced aggression. Two distinct domestication hypotheses posit developmental explanations, involving hypoplasia of embryonic neural crest cells (NCCs), and declining androgen influence, respectively. Here, I assess the operation and potential interactions between these two mechanisms and consider their role in enhanced human adaptation to a cooperative sociocultural niche.

**Methods:** I provide a review and synthesis of related literature with a focus on physiological mechanisms effecting domesticated reductions in masculinity and sexual dimorphism. Further, I examine pre-historic modes of socio-sexual selection likely to drive human self-domestication via reduced aggression and masculinity.

**Results:** I find pluripotent NCCs provide progenitors for a wide range of vertebrate masculine features, acting as regular targets for sexually driven evolutionary change; suggesting domesticated hypoplasia of NCC-derived tissues would be sufficient to explain declines in masculine traits and features. However, lineage specific androgen receptor variability likely moderates these NCC-based effects.

**Conclusions:** These findings extend theorised mechanisms driving noted physiological, morphological, and behavioural changes thought to indicate enhanced sociability and human and self-domestication. Multiple current explanations for human sociability are consistent with physiological domestication under socio-sexual selection favouring dampened masculine physiology and behaviour as adaptations to an enhanced sociocultural niche. The analysis highlights multiple avenues for further investigation.

## 1. Introduction

Controlled breeding experiments, in foxes, mink, chickens, and rats demonstrate that a recognised syndrome of correlated traits common to many domesticated lineages (hereafter ‘domestication syndrome’) tends to emerge in response to sustained selection against reactive aggression, or ‘fight or flight’ type autonomic stress responses (Agnvall, Bélteky, Katajamaa, & Jensen, 2017; Albert et al., 2008; Kharlamova, Faleev, & Trapezov, 2000; Singh et al., 2017; Trut, Oskina, & Kharlamova, 2009). This work supports observational research showing a range of associated behavioural, physiological, and morphological differences in domesticated populations when compared to wild relatives or ancestors (for review see: Hemmer, 1990; Leach, 2003). Recognised traits of domestication syndrome include: diminished aggression and flight distance (behavioural docility); reduced sexual dimorphism; smaller body sizes; less robust skeletons; smaller brains and crania; reduced prognathism; smaller teeth; floppy ears; altered pigmentation; and changed patterns of maturation, oestrus cycling, and fertility (Hemmer, 1990; Kruska, 1988; Leach, 2003; Sánchez-Villagra, Geiger, & Schneider, 2016; Trut et al., 2009; Wilkins, Wrangham, & Tecumseh Fitch, 2014; Zeder, 2012; Zeuner, 1963).

Modern demonstration that selection for tame behaviour triggers this syndrome of unselected traits implies similar changes during Neolithic animal domestication—or pre-Neolithic in the case of dogs, see: Morey and Jeger (2015)—also involved selection for reduced aggressive reactivity (Agnvall et al., 2017; Bidau, 2009; Trut et al., 2009). In this regard, previous authors have argued early domesticators would reflexively have culled aggressive male animals for simple reasons of safety, suggesting this activity altered domesticated populations in various ways (Clutton-Brock, 1984; Helmer, Goucherin, Monchot, Peters, & Sana Segui, 2002; Kruska, 1988; Zeder, 2012). Similarly, in relation to ancient humans themselves, Cieri et al. (2014) suggest pre-historic decline in the craniofacial masculinity of *Homo sapiens* over the past 300,000 years indicates socio-sexual selection against typically-masculine reactive aggression in our own species. Based on a range of documented changes several authors infer a process of ‘human self-domestication’ which is thought to have promoted increased levels of sociability, thereby facilitating language development, group collaboration, knowledge transfer, and technological accumulation (Benítez-Burraco & Kempe, 2018; Cieri et al., 2014; Groves, 1999; Hare, 2017; Thomas & Kirby, 2018; Wrangham, 2019a, 2019b).

According to Cieri et al. (2014) the physiology of human self-domestication involved either reduced in-utero testosterone exposure, or dampened expression of androgen receptors within relevant cells and tissues. In apparent contrast, however, concurrent research into domestication syndrome in non-human animals proposes hypoplasia of neural crest cell (NCC) derived tissues as the defining characteristic of most domesticated traits (Wilkins et al., 2014). Whilst ostensibly diagnosing an alternative mechanism, this latter observation effectively supports Cieri et al.’s (2014) methodology and findings since vertebrate craniofacial morphology is substantially composed from embryonic NCCs (Mishina & Snider, 2014). In fact, as the present article documents (Section 3), many conspicuous secondary sexual features of male vertebrates derive from this cell lineage.

Here, I examine both the NCC hypoplasia and androgen hypotheses of domestication and human self-domestication and consider their implications for human social evolutionary research. I begin by presenting existing findings and perspectives on domestication syndrome and describing the NCC-hypoplasia hypothesis as proposed by Wilkins et al. (2014). I then discuss ancient reductions in masculine traits among domesticated animal lineages and highlight repeated developmental linkages between embryonic NCCs and vertebrate male secondary sexual features. These associations imply that selection against masculine-typical traits—often including elevated reactive aggression—should influence NCC-derived tissues more broadly, thereby promoting domestication syndrome within a given lineage. I proceed to describe NCC contributions to masculine features in humans and discuss pre-historic reduction in human masculinity as an indicator of human self-domestication.

Following this, I consider Cieri et al.’s (2014) proposal that domestication syndrome occurs via altered androgen hormone influence. On balance, given repeated associations between NCCs and masculine vertebrate traits, I argue hypoplasia of NCC-derived tissues would be sufficient to cause domesticated reductions in masculinity and sexual dimorphism; however, androgen receptor placement conspicuously moderates the growth and development of these masculine features. I consider the likely implications of these mechanisms with reference to sociosexual selection for self-domestication in human evolution, finding physiological domestication is consistent with multiple current theories concerning human sociability and our adaptation to an increasingly complex sociocultural niche. In light of this analysis, I discuss avenues for expanded empirical investigation of the human self-domestication hypothesis.

## 2. NCCs and domestication syndrome

The term ‘domestication syndrome’ describes a diverse suite of traits which domesticated animals tend to share in common when compared to wild relatives or ancestors (Hemmer, 1990; Leach, 2003, 2007; Sánchez-Villagra et al., 2016; Trut, 1999; Wilkins et al., 2014; Zeder, 2015). This phenomenon has been a subject of scientific interest and speculation since Darwin’s (1859, 1868) early observations of ‘correlated variation’ in animal domesticates. Symptoms of domestication syndrome include: reduced reactive aggression (behavioural docility); less sexual dimorphism; reduced prognathism; skeletal gracility; smaller teeth; smaller crania and brain size; diminished size and function of sensory organs; changes in pigmentation; heterochrony (shifted rates and timing throughout ontogeny—typically occurring as trait paedomorphisms); changed reproductive behaviour and seasonality; and earlier onset of fertility (Hemmer, 1990; Kruska, 1988; Leach, 2003; Sánchez-Villagra et al., 2016; Trut, 1999; Wilkins et al., 2014; Zeder, 2008, 2015). Based on the observation that many of these effects involve features derived directly from embryonic NCCs, Wilkins et al. (2014) proposed that domestication syndrome results from the heritable hypoplasia of NCC-derived tissues and structures.

NCCs are a transient and pluripotent cell lineage involved in the formation of the vertebrate neural tube during embryonic development (Fig.1). Following normal involution of this tube, these cells migrate on predetermined pathways throughout the developing embryo, providing cellular progenitors for various neural, endocrine, pigment, cardiac, skeletal, and dental cells and tissues (Gilbert, 2010; Le Douarin & Kalcheim, 1999; Schoenwolf, Bleyl, Brauer, & Francis-West, 2008). They contribute to the production and patterning of multiple structures, including: the bones, connective tissues, muscles, and dermis of the craniofacial region; the bones of the middle-ear; the teeth; the hyoid and larynx; pigmented chromatophores (including melanocytes) of vertebrate skin, hair, and feathers; various glands including the thymus, the anterior pituitary, the adrenal medulla and associated autonomic nerve structures; as well as parts of the vertebrate heart (Gilbert, 2010; Le Douarin & Kalcheim, 1999; Schoenwolf et al., 2008; Ueharu et al., 2017). Due to their diverse contributions, NCCs exert a considerable influence on the morphology, physiology and behaviour of all vertebrate taxa.

**Fig. 1.**
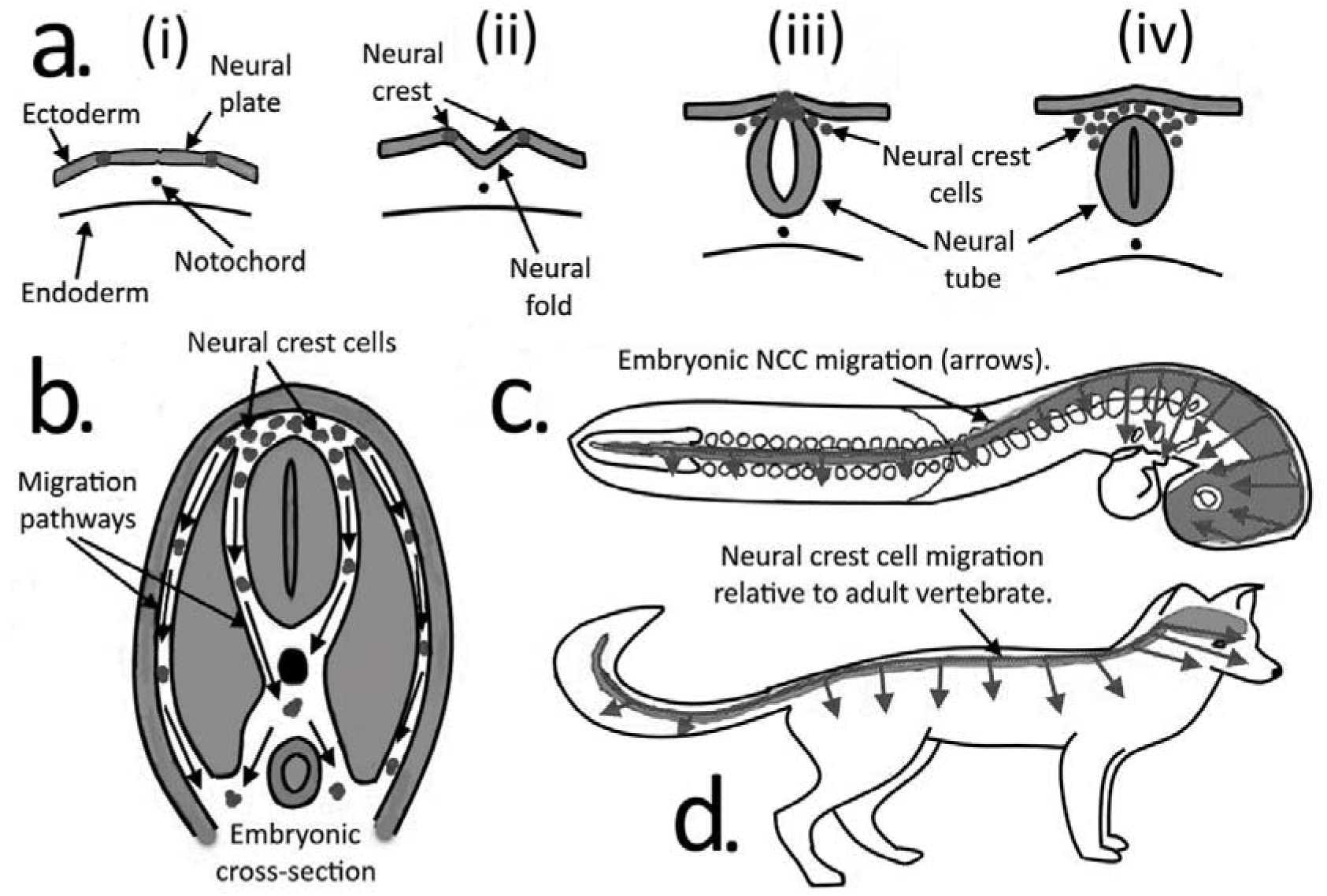
Embryonic NCC function and migration pathways. (**a**) (i) through (iv): Formation of the neural tube: converging edges of the involuted neural fold are fused by NCCs which subsequently migrate throughout the developing embryo. (**b**): Embryonic cross-section indicating migration of NCCs following neural tube formation. (**c**): Idealised NCC dispersal to various regions in a developing vertebrate embryo. (**d**): The dispersal pathways shown in (c) relative to adult body plan of mammalian vertebrate. (image adapted from: Le Douarin, Creuzet, Couly, & Dupin, 2004; A. Liu & Niswander, 2005; Wilkins et al., 2014; Gilbert, 2010)

Results from long-running experiments using captive silver foxes (*Vulpes vulpes*) demonstrate that sustained breeding selection against aggressive stress reactivity provokes a range of unselected domestication-typical traits (‘domestication syndrome’) within a given lineage (Trut, 1999; Trut et al., 2009). Similar effects are documented in mink (Kharlamova et al., 2000; Kulikov et al., 2016), in rats (Albert et al., 2008; Singh et al., 2017), in chickens (Agnvall et al., 2017; Jensen, 2006), and in semi-domesticated mice (Geiger, Sánchez-Villagra, & Lindholm, 2018). Rather than repeated random mutation of genes controlling each commonly altered feature across multiple domesticated lineages and taxa, many domesticated traits are proposed to result from shared developmental shifts (Belyaev, 1979; Fallahsharoudi et al., 2015; Osadchuk, 1998; Trut et al., 2009). In accord with these expectations, both the NCC hypoplasia (Wilkins et al., 2014) and androgen (Cieri et al., 2014) hypotheses posit broadly-influential mechanisms of ontogenic change.

At present, modified NCC functioning provides the most widely supported explanation for the observed link between selection for reduced aggression and the wider syndrome of domesticated traits (Geiger et al., 2018; Montague et al., 2014; Pendleton et al., 2018; Wilkins, 2017; Wilkins et al., 2014; Zeder, 2017). Wilkins et al. (2014) proposed that hypoplasia of the NCC-derived adrenal medulla dampens behavioural reactivity prompted by the Hypothalamic-Pituitary-Adrenal (HPA) axis. Elsewhere, Wrangham (2019b) suggests known NCC influence upon the patterning of forebrain development (Creuzet, 2009) explains smaller limbic systems observed in domesticated lineages (Kruska, 1988, 2005). In addition, NCC contributions to the pituitary (Ueharu et al., 2017) imply that hypoplasia of this organ might also dampen HPA reactivity. In effect, therefore, selection against reactive aggression is thought to alter the development of autonomic nervous system components via heritable changes to NCC functioning, and, since NCCs provide progenitors for a range of other features, this promotes the broader suite of correlated changes seen in domestication syndrome (Wilkins et al., 2014) (Fig.2).

**Fig. 2.**
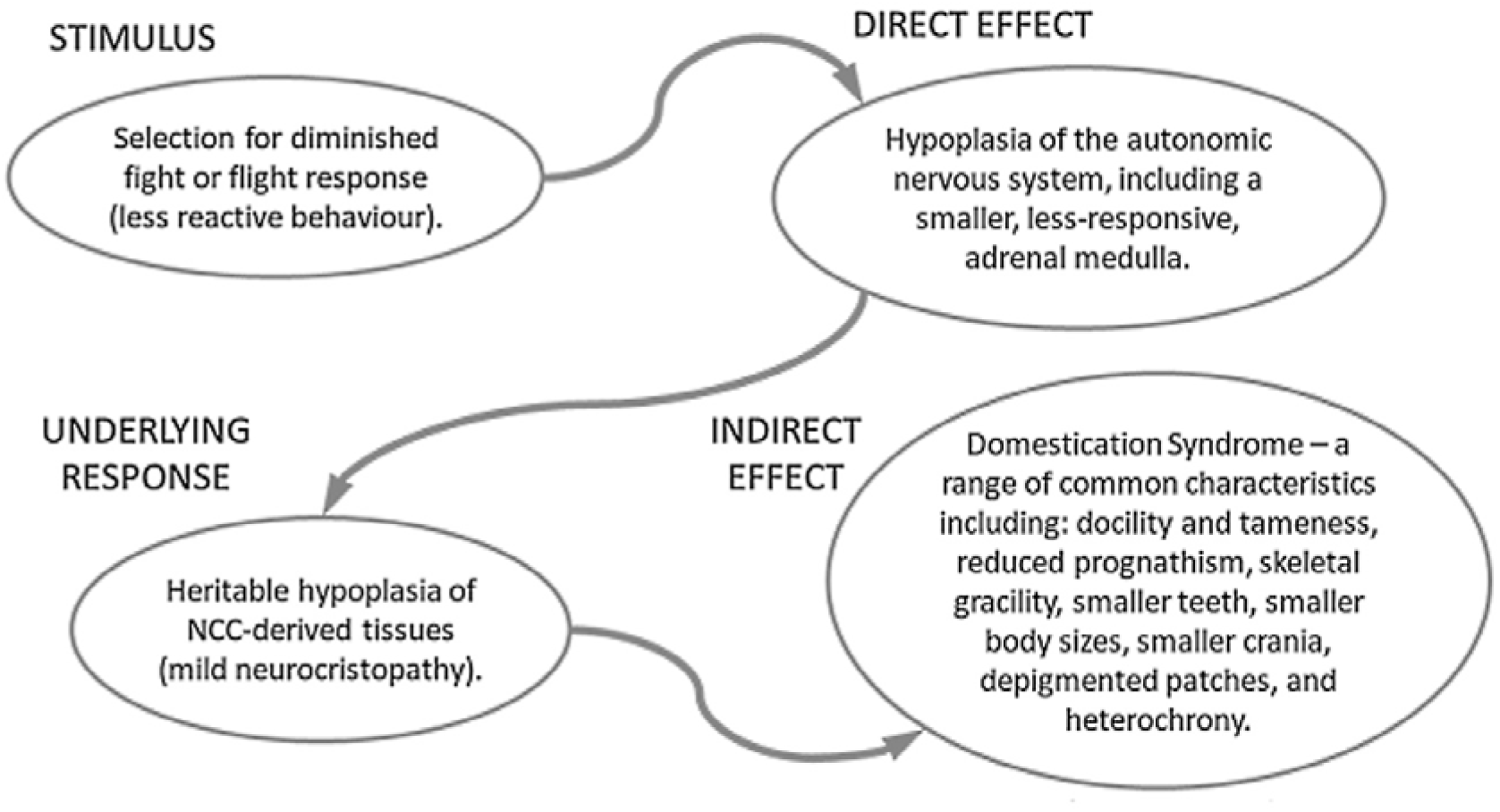
Proposed chain of influence in vertebrate domestication from Stimulus (selection for less reactive or aggressive behaviour) to Indirect Effect (domestication syndrome) (Wilkins et al., 2014)

Wilkins et al. (2014) suggest domesticated hypoplasia of NCC-derived tissues occurs via either: (1) a reduction in the original number of embryonic NCCs at the neural tube; (2) a dampening of NCC migration leading to fewer cells at their final destinations; or (3) a decline in in-situ proliferation or growth at those destinations. On balance, they lean towards the migration hypothesis based largely on the observation that common piebald patterns of depigmentation in domesticates occur at sites relatively distant from the embryonic neural tube (feet, belly, tail-tip, and forehead), implying constrained movement of NCCs from their point of origin (Wilkins et al., 2014). It is assumed that the same dampened migratory capacity drives apparent hypoplasia in other NCC-derived tissues and structures. Genetic research has identified a range of signals and regulators acting in complex gene regulatory networks to control NCC specification, delamination, migration, and differentiation into their derivative tissues (Simões-Costa & Bronner, 2015). Study of piebald depigmentation points to KIT gene mutations which dampen NCC migration ability, causing an absence of melanocytes in zones distal to the neural tube (Grichnik, 2006; Oiso, Fukai, Kawada, & Suzuki, 2013).

## 3. Domestication and masculine traits

Charles Darwin (1859, 1868) originally posited a period of ‘unconscious selection’ which must logically have preceded any intentional human cultivation for desirable traits in captive animals. Several subsequent authors suggest early domesticators would reflexively have eliminated particularly aggressive male animals for simple reasons of safety and self-preservation (Clutton-Brock, 1984; Helmer et al., 2002; Kruska, 1988; Zeder, 2012) thereby enacting a persistent, if initially ‘unconscious’, selection against aggressive reactivity. Low y-DNA diversity in many modern lineages—including: dogs, horses, cattle and sheep (Lippold et al., 2011)—supports this expectation of relatively higher selection on early male domesticates. Higher diversity in mtDNA, combined with observations of traditional husbandry practices involving culling and castration of juvenile males (Marshall, Dobney, Denham, & Capriles, 2014; Zohary, Tchernov, & Horwitz, 1998) add further weight to this hypothesis. Incidentally, similar sex-biased behavioural selection was also present in the Russian fox domestication experiment which started with 100 females and 30 males and in subsequent generations, on average, kept only 3% of the most sociable males as breeders, in comparison to 8-10% of females (Trut, Plyusnina, & Oskina, 2004).

Accepting that reflexive male-biased behavioural selection (a bias *against* aggressive males) was a recurring factor throughout the pre-history of major domesticates, it is not surprising that male phenotypes are typically more substantially altered than those of females when compared to non-domesticated ancestors or relatives (Cieri et al., 2014; Sánchez-Villagra et al., 2016; Zohary et al., 1998). For example, whilst the majority of wild mammals show males with larger average body size than females (Blanckenhorn, 2005; Rensch, 1950), declines in sexual body size difference (with males becoming relatively smaller) are a recurring feature under domestication (Hemmer, 1990; Polák & Frynta, 2009; Sánchez-Villagra et al., 2016; Zohary et al., 1998). In fact, this effect provides the clearest indicator of domestication in early archaeological deposits (Helmer et al., 2002; Zeder, 2008, 2012). Size reduction, or complete loss, of horns (which are larger and occur more often in male bovids) are a further example of reduced masculinity and sexual dimorphism in lines of domesticated sheep, goats, and cattle (Zeder, 2008, 2012; Zeuner, 1963; Zohary et al., 1998). In addition, experimentally domesticated foxes (Trut et al., 2009, 2004), and mink (Kharlamova et al., 2000), show male-biased decline in prognathism (the projection of masticatory skeleton in relation to the crania), leading to male craniofacial ‘feminisation’ and less sexual difference. Similar reductions in prognathism are a feature of most domesticated lineages (Leach, 2003; Morey & Jeger, 2015; Zeuner, 1963), although whether this has consistently affected males more than females is unknown.

If domestication syndrome, and the reduced sexual dimorphism associated with it, results primarily from altered NCC functioning as proposed by Wilkins et al. (2014), we should logically expect some physiological link between these cells and the features or physiology that causes various sexual differences. In addition, since declining sexual dimorphism occurs primarily via diminution of male-typical traits, this linkage should especially influence the secondary sexual features of males. Such a relationship would be particularly apparent in features directly derived from NCC progenitors since these should tend towards correlated hypoplasia under domestication. However, NCC contribution to the vertebrate pituitary and its hormone-secreting cells (Ueharu et al., 2017) suggest NCC hypoplasia might affect sexual differences more broadly via changed endocrine signalling regimes. Recent investigations support this latter expectation, suggesting altered pituitary function drives important shifts under domestication (Fallahshahroudi, Løtvedt, Bélteky, Altimiras, & Jensen, 2018). Pituitary signalling affects a range of developmentally relevant processes; directing the release of hormones including: adrenaline, growth hormone, thyroid hormone, cortisone, and testosterone (Gilbert, 2010; Schoenwolf et al., 2008). Overall, therefore, generalised hypoplasia of NCC-derived tissues might influence domesticated ontogeny via two primary modes: either directly, following reduced cellular contributions to a range of NCC-derived features; or indirectly, through altered pituitary development, affecting the rate and timing of endocrine signalling throughout ontogeny.

In direct support of this latter effect, heterochronic developmental shifts are a consistent feature of the domestication syndrome—with changed timing of sexual maturity being especially common (Belyaev, 1979; Clutton-Brock, 1984; Hemmer, 1990; Jensen, 2006; Trut, 1999; Wilkins et al., 2014; Zeder, 2012, 2015). Although debate continues as to whether domesticated heterochrony reliably leads to paedomorphism (adult retention of ancestral juvenile traits) (Evin et al., 2017), shifts in maturation would almost certainly influence sexual differences since these regularly emerge via sex-specific developmental rates and timing (Humphrey, 1998; Leigh, 1992; Plavcan, 2001). For instance, larger male body size is not a ‘feature’ composed from NCC-derived tissues, therefore, domesticated reduction in sexual size difference cannot follow directly from NCC-hypoplasia; however, hypoplasia of NCC-derived pituitary cells can explain these changes via altered endocrine regimes. As a relevant aside, mean body size for a given lineage also tends to reduce under domestication (Clutton-Brock, 1999; Hemmer, 1990; Sánchez-Villagra et al., 2016) and is also substantially moderated by endocrine signalling throughout development.

Beyond these developmental difference, many conspicuous sexual dimorphisms involve masculine features directly derived from NCC progenitors (Table 1). In such cases, a link between NCCs and a given feature is inherently obvious. For example, vertebrate teeth derive from NCCs, are regularly used for fighting and aggressive display, and are often highly sexually dimorphic (Andersson, 1994; Emlen, 2008; B. K. Hall, 2000; Le Douarin & Kalcheim, 1999). Masculine features are also regularly composed from elements of the NCC-derived craniofacial skeleton. In primates, for example, these elements may include notably dimorphic brow ridges, cheek flanges, and sagittal crests (Andersson, 1994; Balolia, Soligo, & Wood, 2017; A. Dixson, Dixson, & Anderson, 2005; Plavcan, 2001). Across vertebrates more widely, conspicuous masculine signalling often occurs in the form of hair, skin, and feather colouration, depending on various NCC-derived chromatophores, including melanocytes (Bagnara & Hadley, 1973; Le Douarin & Kalcheim, 1999). These are predominantly darker, or more intensely colourful, in males (Andersson, 1994; Darwin, 1871). Different forms of acoustic signalling provide another common masculine trait (Andersson, 1994), and are typically reliant on NCC-derived structures such as neck muscles, the hyoid, the larynx, and the syrinx.

**Table 1.** Common male vertebrate secondary sexual traits and their links to NCCs

| Male secondary | Taxonomic examples | Role of NCCs |
| --- | --- | --- |

| traits |  |  |
| --- | --- | --- |
| **Craniofacial morphology** | Brow ridges, sagittal crests, cheek flanges and other craniofacial features in primates. | NCCs form the entire vertebrate craniofacial region.<sup>1-4</sup> |
| **Horns, antlers and other headgear.** | Horns, pronghorns, antlers, and ossicones in Ungulates. | NCCs thought to provide cellular progenitors for antlers.<sup>5,6</sup> Implicated in development of horns, pronghorns, and ossicones by forming the dermis and connective tissues from which they emerge, as well as frontal bones to which they attach.<sup>3,7</sup> |
| **Larger teeth.** | Tooth (especially canine) differences common across multiple taxa. | NCCs provide cellular progenitors for all tooth odontoblasts and papillae.<sup>3,7,8</sup> |
| **Pigmented and structural colourations.** | Pigmented and structurally derived feather, skin, and hair colourations in numerous taxa. | NCCs produce chromatophores which create or support a range of vertebrate pigmentations and cellular iridescence in skin, hair, and feathers.<sup>7,9</sup> |
| **Vocal signalling.** | Masculine vocalisations across numerous vertebrate taxa. | NCCs provide progenitor cells for the hyoid, larynx, syrinx, and associated neck and throat muscles.<sup>1,11</sup> |
| **Specific sexual behaviours.** | Elevated male reproductive effort and aggression in multiple taxa. | NCCs influence the autonomic nervous system and testosterone production via adrenal medulla<sup>3,8</sup> and pituitary.<sup>12</sup> NCC patterning of forebrain development may affect limbic components.<sup>13</sup> |
| **Physiological differences.** | Sexual differences of the vertebrate immune system. | NCCs implicated in immune system function via contributions to the vertebrate thymus and bone marrow<sup>7,14</sup> |

Notes: <sup>1</sup>(Bhatt, Diaz, & Trainor, 2013); <sup>2</sup>(Cordero et al., 2011); <sup>3</sup>(Santagati & Rijli, 2003); <sup>4</sup>(Schoenwolf et al., 2008); <sup>5</sup>(Davis, Brakora, & Lee, 2011); <sup>6</sup>(Kierdorf & Kierdorf, 2010); <sup>7</sup>(Le Douarin & Kalcheim, 1999); <sup>8</sup>(Le Douarin et al., 2004); <sup>9</sup>(Bagnara & Hadley, 1973); <sup>10</sup>(Ligon, Thornhill, Zuk, & Johnson, 1990); <sup>11</sup>(Mishina & Snider, 2014); <sup>12</sup>(Ueharu et al., 2017); <sup>13</sup>(Wrangham, 2019b); <sup>14</sup>(B. K. Hall, 2000).

In summary, NCCs are a pluripotent cellular lineage with the capacity to form a wide variety of cells, tissues, organs, and structures. The exceptional diversity of male secondary sexual traits derived from this lineage suggests repeated involvement in processes of sexual recognition, male-male competition, and female choice. As discussed, noted NCC contributions to the vertebrate pituitary (Ueharu et al., 2017), combined with developmental shifts commonly seen under domestication, imply a further influence on sexual differences from NCC hypoplasia via moderated endocrine regimes. These cross-phylogenetic contributions to the expression of divergent sexual characteristics suggest NCCs play an important role as regular targets of selection, and facilitators of taxonomic differentiation.

## 4. NCCs and human masculinity

Multiple previously designated features of human masculinity also derive directly from embryonic NCCs (Table 2). As in other vertebrates, these cells provide progenitors for the entire human craniofacial region, including: the frontal bone and associated brow ridges, the mandible and maxilla, the nasal bone and cartilage, and the zygomatic arches, as well as influencing the patterning of overlying facial muscles (Gilbert, 2010; Knight & Schilling, 2013). As such, cranial NCCs form the cellular foundations for all proposed indicators of human facial masculinity, including the facial width to height ratio widely discussed within human behavioural and sexual selection literature (e.g. see Carré, McCormick, & Mondloch, 2009; Feinberg, DeBruine, Jones, & Little, 2008; Lefevre, Lewis, Perrett, & Penke, 2013; Mitteroecker, Windhager, Müller, & Schaefer, 2015), and used by Cieri et al. (2014) as an indicator of masculinity and self-domestication in ancient humans.

**Table 2.** Masculine human traits and their relation to NCCs

| Noted masculine traits in humans | Role of NCCs |
| --- | --- |
| **Robust facial morphology and proportions including brow-ridges and chin.**<sup>1–8</sup> | NCCs form all of the facial skeleton, including the frontal bone, mandible, maxilla, zygomatics, and nasal structures, as well as associated muscle, cartilage, and connective tissue.<sup>9–11</sup> |
| **Large teeth.**<sup>12,13</sup> | NCCs provide progenitors for all vertebrate teeth.<sup>14–16</sup> |
| **Thickened facial hairs (beards).**<sup>17–19</sup> | NCCs produce the dermis of the face and neck, including associated hair follicles.<sup>8,20,21</sup> |
| **Adam's apple and lower vocal pitch.**<sup>22–24</sup> | NCCs provide progenitor cells for the hyoid, larynx, and associated neck muscles.<sup>6,25</sup> |
| **Shoulder width and shoulder to hip ratio.**<sup>26,27</sup> | NCCs provide progenitors for parts of the shoulder girdle and connective tissues and influence developmental patterning of associated muscles.<sup>28–31</sup> |
| **Elevated status-striving, interpersonal aggression, and violence.**<sup>32, 33</sup> | NCC contributions to pituitary, adrenal medulla, and limbic patterning imply influences on temperament and behavioural responses via HPA reactivity and testosterone release.<sup>34–36</sup> |

Notes: <sup>1</sup>(Carré et al., 2009); <sup>2</sup>(Carrier & Morgan, 2015); <sup>3</sup>(Hodges-Simeon, Sobraske, Samore, Gurven, & Gaulin, 2016); <sup>4</sup>(Lefevre et al., 2013); <sup>5</sup>(Mitteroecker et al., 2015); <sup>6</sup>(Cieri et al., 2014); <sup>7</sup>(B. J. Dixon, 2016); <sup>8</sup>(Shearer, Sholts, Garvin, & Wärmländer, 2012); <sup>9</sup>(Mishina & Snider, 2014); <sup>10</sup>(Santagati & Rijli, 2003); <sup>11</sup>(Schoenwolf et al., 2008); <sup>12</sup>(Coquerelle et al., 2011); <sup>13</sup>(Schwartz & Dean, 2005); <sup>14</sup>(Gilbert, 2010); <sup>15</sup>(B. K. Hall, 2008); <sup>16</sup>(J. E. Hall, 2010); <sup>17</sup>(Craig, Nelson, & Dixon, 2019); <sup>18</sup>(B. J. Dixon & Brooks, 2013); <sup>19</sup>(B. J. Dixon, Lee, Sherlock, & Talamas, 2017); <sup>20</sup>(Jinno et al., 2010); <sup>21</sup>(Krause et al., 2014); <sup>22</sup>(Feinberg et al., 2008); <sup>23</sup>(Puts, Gaulin, & Verdolini, 2006); <sup>24</sup>(Puts et al., 2016); <sup>25</sup>(Bhatt et al., 2013); <sup>26</sup>(Lee, 2015); <sup>27</sup>(Windhager, Schaefer, & Fink, 2011); <sup>28</sup>(Matsuoka et al., 2005); <sup>29</sup>(McGonnell, McKay, & Graham, 2001); <sup>30</sup>(Noden, 1986); <sup>31</sup>(Valasek et al., 2010); <sup>32</sup>(M. Wilson & Daly, 1985); <sup>33</sup>(Archer, 2009); <sup>34</sup>(Ueharu et al., 2017); <sup>35</sup>(Schoenwolf et al., 2008); <sup>36</sup>(Creuzet, 2009).

NCCs also compose the dermis of the face and neck, including associated hair follicles, thus providing progenitor tissues for the human beard (Jinno et al., 2010; Krause et al., 2014; Schoenwolf et al., 2008)—a highly sexually dimorphic trait and another focus of sexual selection research (B. J. Dixson & Brooks, 2013; B. J. Dixson, Rantala, Melo, & Brooks, 2017). Further, they compose the human larynx and hyoid (Schoenwolf et al., 2008), substantially influencing human vocal qualities, including voice pitch—another focus of study regards human masculinity (Feinberg et al., 2008; Puts et al., 2016; Puts, Jones, & DeBruine, 2012). NCC contributions to the development of the vertebrate shoulder region influence its relative size and robusticity by forming sections of the clavicle and scapula bones, providing muscle connection sites, and directing developmental patterning in major neck and shoulder muscles (Ericsson, Knight, & Johanson, 2013; Matsuoka et al., 2005; McGonnell et al., 2001; Noden, 1986; Valasek et al., 2010). Assuming no associated change in the (non-NCC-derived) pelvis, this implies NCCs might influence shoulder width, and the ‘shoulder-to-hip-ratio’; both of which are proposed as indicators of masculinity and targets of sexual selection (Lee, Brooks, Potter, & Zietsch, 2015; Windhager et al., 2011).

Finally, NCCs are also substantially implicated in the physiology of masculine behaviours. As mentioned, their contributions to the adrenal medulla (Schoenwolf et al., 2008; Wilkins et al., 2014), the pituitary (Ueharu et al., 2017), and, possibly, the patterning of limbic structures (Wrangham, 2019b) testifies to substantial involvement in autonomic stress reactivity initiated via the HPA axis. NCC contributions to the pituitary imply a further behavioural influence via patterns of endocrine signalling; especially in its control of luteinizing hormone which stimulates testosterone release from male testes. Testosterone has been associated with male sex-drive and aggressive behaviour across a range of species, including in humans (Harding, 1983; Harris, 1999; Lincoln, 1989); however, recent research supports a less direct association with male aggression, implying testosterone’s main effect is to promote status striving (Carré & Archer, 2018) which need not involve overtly aggressive -actions. In support of this view, testosterone has been associated with both aggressive *and pro-social* (cooperative and sharing) behaviours in humans; the latter particularly in situations where generosity might enhance social status (Dreher et al., 2016).

## 5. Signs of domestication in recent humans

From the Mid-Pleistocene, human evolution has involved substantial decline in several sexually dimorphic features, suggesting reductions in relative masculinity and a shift towards a more gracile (‘feminised’) morphology (Brace & Ryan, 1980; Cieri et al., 2014; Frayer, 1980; Frayer & Wolpoff, 1985; Hill, Bailey, & Puts, 2017; Ruff, Trinkaus, & Holliday, 1997). These trends are consistent with changes seen in domestication syndrome, leading multiple authors to suggest that *Homo sapiens* has undergone a physiological process of self-domestication (Cieri et al., 2014; Franciscus, Maddux, & Schmidt, 2013; Groves, 1999; Hare, 2017; Leach, 2003; Wrangham, 2018, 2019b, 2019a). Perhaps the most indicative shifts include declines in body size and body size sexual dimorphism (Frayer, 1980; Frayer & Wolpoff, 1985; Gallagher, 2013; Hill et al., 2017; Ruff, 2002; Ruff et al., 1997; Ryan & Shaw, 2015), although teeth and cranial size and shape also show comparatively less sexual difference in contemporary populations (Brace & Ryan, 1980; Frayer & Wolpoff, 1985). In addition, there has been marked reduction in skeletal robusticity (Ryan & Shaw, 2015), including in the cranial and facial skeleton (Carrier & Morgan, 2015; Cieri et al., 2014; Frayer, 1980), along with reductions in facial projection and prognathism (Carrier & Morgan, 2015; Cieri et al., 2014; Lieberman, 1998, 2011; Polychronis & Halazonetis, 2014) and decline in mean tooth size, with increased rates of tooth agenesis (Brace, Rosenberg, & Hunt, 1987; Calcagno & Gibson, 1988; Polychronis & Halazonetis, 2014). Finally, in contrast with longstanding evolutionary trends toward greater hominin encephalisation, recent humans show a marked decline in cranial capacity compared to earlier *Homo sapiens* (Balzeau et al., 2012; Henneberg, 1988; Hublin Jean-Jacques, Neubauer Simon, & Gunz Philipp, 2015; Lieberman, 2011; C. Liu et al., 2014; McHenry, 1994; Ruff et al., 1997; Wiercinski, 1979).

Despite reduced sexual differences in contemporary human populations, many traits remain sexually bimodal today; with men being physically larger and more robust on average (Coquerelle et al., 2011; Frayer & Wolpoff, 1985; Gaulin & Boster, 1992; Pettenati-Soubayroux, Signoli, & Dutour, 2002; Wells, 2012). Worldwide, stature sexual dimorphism varies around a mean of 7% larger males (Gaulin & Boster, 1992; Gleeson & Kushnick, 2018; Ralls, 1976). In addition, male craniofacial measurements (5-9%) and tooth size (6.5%) are also larger on average (Schwartz & Dean, 2005; Ursi, Trotman, McNamara, & Behrents, 1993). Men show higher incidence of hyperdontia (supernumerary teeth) and megadontia, whereas women more often show hypodontia (tooth agenesis) and microdontia (Anderson, Thompson, & Popovich, 1975; Brook, 1984; Polder, Hof, Linden, & Kuijpers□Jagtman, 2004). These continuing sexual differences suggest associated pre-historic reduction in body and craniofacial size and robusticity, along with smaller tooth sizes and increased agenesis, all constitute a relative ‘feminisation’ of the phenotypic mean.

Behaviourally, relative to women, contemporary males show elevated propensity for interpersonal aggression and violence, both internationally and cross-culturally (Fry & Söderberg, 2013; UNODC, 2013). These sex-linked predispositions are well-documented and widely discussed among evolutionarily-focussed authors (e.g. see Archer, 2009; Buss & Shackelford, 1997; Georgiev, Klimczuk, Traficonte, & Maestripieri, 2013; M. Wilson & Daly, 1985). Noted sexual differences (both behavioural and morphological) are regularly attributed to divergent sexual selection; especially reproductive competition as physical conflict among males (Carrier & Morgan, 2015; Fink, Weege, Manning, & Trivers, 2014; Hill et al., 2017; Scott, Clark, Boothroyd, & Penton-Voak, 2013; Sell, Hone, & Pound, 2012). This explanation is theoretically compelling, and has been explored empirically via comparisons of sexual dimorphism under monogamy and polygyny across different species, and in different human societies—the latter with mixed results. (Alexander, Hoogland, Howard, Noonan, & Sherman, 1979; A. F. Dixson, 2012; Gaulin & Boster, 1992). Despite widespread expectations of strong association between sexual dimorphism and polygyny, sexual size difference in non-human primates better reflects simple frequencies of agonistic interaction (Plavcan, 2012; Plavcan & van Schaik, 1997). This observation suggests sexual dimorphism indicates levels of aggression and male social intolerance, rather than polygyny per se. As such, decline in human sexual dimorphism would be associated with reductions in agonistic social interaction, irrespective of social or sexual arrangements.

In support of this expectation, noted pre-historic decline in morphological masculinity and sexual dimorphism coincides with an increasing cultural and technological complexity, widely thought to result from elevated sociability and collaboration (Boehm, 2014; Burkart et al., 2014; Cieri et al., 2014; Flinn, Geary, & Ward, 2005; Hare, 2017; Hawkes, 2013; Henrich, 2017; Hrdy, 2009; Sterelny, 2011). Whilst multiple factors have been proposed to explain these apparent socio-cognitive shifts, existing inference tends to avoid analysis of biophysical mechanisms that might affect human social behaviour. In distinct contrast, however, the explicit proposition at the core of the human self-domestication hypothesis is that documented socio-cultural change and documented morphological change are associated via the same physiological process observed in domestication syndrome (Cieri et al., 2014; Hare, 2017; Wrangham, 2018, 2019b, 2019a); an effect repeatedly demonstrated to emerge under selection for reduced behavioural reactivity and aggression (Albert et al., 2008; Jensen, 2006; Kharlamova et al., 2000; Trut, 1999; Trut et al., 2009). In essence, therefore, the human self-domestication hypothesis offers a compelling physiological foundation for several existing lines of inference regards the role and causes of enhanced human sociability and collaboration. This is not an attempt to assert a form of biological determinism in the evolution of human behaviour. Instead, it identifies a suite of morphological, behavioural, and physiological evidence which can help to explain why (and, especially, *how*) human social interaction is the preeminent force behind our uniquely successful cultural and technological adaptive strategies. It explicitly asserts that human sociability and cultural development shaped the biophysical evolution of our species to suit an increasingly complex and cooperative sociocultural niche.

## 6. Considering the androgen hypothesis

Having described widespread NCC contributions to masculine vertebrate morphology and behaviour, and their theorised role in the physiology of domestication, I now turn to consider Cieri et al.’s (2014) androgen hypothesis. This proposal was specifically formulated in relation to *human* self-domestication over the past 200,000 years, but by implication describes mechanisms common to domestication generally. The hypothesis suggests that past reduction in *Homo sapiens*’ craniofacial masculinity is evidence of self-domestication caused either by lower levels of circulating androgens, or by changes in androgen receptor function (Cieri et al., 2014). Sánchez Villagra and van Schaik (2019) recently discussed this proposal (and several issues raised in connection with it), but did not explicitly distinguish it from the NCC hypoplasia hypothesis. Here, I consider these two hypotheses separately since neither Cieri et al. (2014), nor Wilkins et al. (2014), provide any link between them.

As is widely apparent, nominally masculine features respond predictably to varying quantities of circulating androgens throughout the lifecycle of a given individual (Bardin & Catterall, 1981; Gilbert, 2010). In male humans, elevated testosterone at puberty is known to promote developmental changes, including: growth in the NCC-derived larynx, leading to: altered vocal pitch (Hollien, Green, & Massey, 1994); thickened facial hairs (beards) from the NCC-derived dermis of the face and neck (Rutberg et al., 2006); an overall increase in structural masculinity of the NCC-derived facial region (Zaidi et al., 2019); as well as other physiological shifts (Gilbert, 2010). Levels of circulating testosterone tend to correlate with facial masculinity in adolescence and adulthood (Lefevre et al., 2013; Marečková et al., 2011) and recent research suggests links between prenatal androgen exposure and the masculinity of human faces at birth (Whitehouse et al., 2015; Zaidi et al., 2019). Similarly, twin studies show influence from intrauterine androgens upon sexual tooth size difference (Ribeiro, Brook, Hughes, Sampson, & Townsend, 2013).

Beyond this comparative evidence, experimental testosterone treatment in boys with unusually delayed puberty has been shown to promote increased upper and total facial height (Verdonck, Gaethofs, Carels, & de Zegher, 1999), demonstrating a direct effect upon NCC-derived craniofacial morphology—particularly, the facial width-to-height ratio, a measurable indicator of facial masculinity (Cieri et al., 2014; Hodges-Simeon et al., 2016; Lefevre et al., 2013). Similarly, administration of anabolic steroids in young rats caused increased length of the craniofacial skeleton, especially the midface, maxilla, and mandible (Barrett & Harris, 1993), whilst, conversely, neonatal and prepubertal castration caused diminished craniofacial growth, producing a shorter anterior length (Verdonck et al., 1998) as commonly seen under domestication syndrome (Trut et al., 2009; Wilkins et al., 2014; Zeuner, 1963).

In accord with Cieri et al.’s (2014) androgen hypothesis, these observations demonstrate androgenic influence across a wide range of masculine features—as would be expected given well-established links between androgens and masculinity. It may appear, therefore, that domesticated declines in masculinity arise simply from reduced circulating androgens. In accord with this possibility, experimentally-domesticated male foxes do show reduced testosterone (Osadchuk, 2001); however, relatively *elevated* testosterone has been reported for several other domesticates, including: zebra finches (Prior et al., 2017); guinea pigs (Künzl & Sachser, 1999; Zipser, Schleking, Kaiser, & Sachser, 2014); and domesticated sheep breeds and crossbreeds—excepting merinos—(Lincoln, Lincoln, & McNeilly, 1990).

Given this inconsistent trend, it seems that simple testosterone reduction cannot be responsible for the shared range of traits seen in domestication syndrome. However, regular reductions in domesticated masculinity and sexual dimorphism, despite occasionally increased testosterone, may support Cieri et al.’s (2014) alternative proposition of altered androgen receptor functioning. Although typically ignored in testosterone-focussed studies, varied expression and placement of androgen receptors are crucial moderators of androgen hormone effects (Matsumoto et al., 2013; Monks & Holmes, 2017). These cellular receptors act as both transcription factors—to enable genomic pathways of androgen influence—as well as promoters of rapid transcription-independent androgen response (Matsumoto et al., 2013; Palvimo, 2012). They moderate growth and development of cells and tissues via hyperplasia or hypertrophy (increased number and size of cells, respectively) (Emerson, 2000; Palvimo, 2012), and are distributed as appropriate to taxonomically specific modes of masculine signalling and sexual behaviour; with somatic constraints enforcing a level of optimisation and trade-offs (Monks & Holmes, 2017). As such, whilst fluctuations in circulating testosterone may induce seasonal or lifecycle shifts in masculine traits (including seasonal behaviours), varied distributions of androgen receptors necessarily target these effects to species-relevant features; emphasising the expression of specific sexual morphologies, vocalisations, colourations, and behaviours.

Widespread androgenic influence upon NCC-derived masculine features might suggest a synthesis of the androgen receptor (Cieri et al., 2014) and NCC (Wilkins et al., 2014) hypotheses, whereby domestication would dampen the functioning of androgen receptors specifically within NCC-derived tissues. According to this proposal, domesticated hypoplasia of NCC-derived tissues would, therefore, emerge via a weakening of normal *hyper*plasic influence from androgens. One observation counting against this speculative proposal, however, is that androgen effects on NCC-derived melanocytes do not easily explain piebald patterns of depigmentation—a recognised neurocristopathy commonly seen under domestication. Whilst androgens are known to influence melanocyte function (M. J. Wilson & Spaziani, 1973), and both androgen levels and receptor locations regularly influence sexual colour differentiation (Slominski, Tobin, Shibahara, & Wortsman, 2004), piebald patterning more closely accords with predictions of dampened NCC migration (Wilkins et al., 2014). Unlike albinism, where melanocytes are present but do not produce melanin, piebaldism involves an absence of NCC-derived melanocytes in affected areas (Pavan & Tilghman, 1994). As mentioned earlier, genetic research has identified various KIT gene mutations that cause piebalding via the influence of KIT ligand (or ‘Stem Cell Factor’), a known facilitator of NCC migration (Alexeev & Yoon, 2006; Grichnik, 2006; Oiso et al., 2013).

Interestingly, female androgen hormones are thought to influence KIT ligand production as part of the reproductive biology of mice, acting via androgen receptors located in granulosa cells of the ovarian follicle (Matsumoto et al., 2013). This example indicates a moderating relationship between androgen receptors and the production of KIT ligand. If a similar mechanism were to operate within fertilised embryos, this would suggest an upstream regulatory function for androgen receptors in early NCC migration, thus directly linking them to piebald patterning—and other NCC migration-related symptoms of domestication syndrome. Having said this, the possibility of such an embryonic mechanism is entirely speculative.

In summary, pluripotent lineages of embryonic NCCs regularly provide cellular progenitors for lineage-specific masculine traits across the vertebrates (see Tables 1 and 2). In addition, varied distributions of androgen receptors moderate the development of these cellular foundations, targeting steroidogenic hypertrophy and hyperplasia to specific cells and tissues thereby emphasising sexually relevant features in a given individual or lineage. Despite conspicuous androgenic effects upon NCC-derived expressions of masculinity, however, common patterns of domesticated piebalding clearly involve dampened NCC migration caused by mutations affecting the functioning of KIT ligand (Alexeev & Yoon, 2006; Oiso et al., 2013). It follows that whilst androgen receptor expression and placement may moderate in-situ NCC-hypoplasia under domestication, this effect is probably less influential as a shared cause of masculinity-decline than that of dampened NCC migration. The possibility that androgen receptors might provide an upstream influence upon the production of KIT ligand, and could, therefore, moderate embryonic NCC migration, requires further exploration.

## 7. Selection against masculinity in humans

Previous research explicitly focussed on human self-domestication has generated three primary hypotheses in regard to sociosexual selection against reactive aggression in humans; these are: (1) social benefits (Cieri et al., 2014; Hare, 2017; Sánchez Villagra & van Schaik, 2019), whereby improved sociability would provide enhanced survival and fitness due to elevated capacity for cooperation and exchange; (2) collaborative execution (Wrangham, 2018, 2019a, 2019b), where sociable group members would conspire to inflict capital punishment on excessively aggressive and dominating individuals; and (3) female mate choice (Cieri et al., 2014; Gleeson & Kushnick, 2018), whereby women preferentially select for less masculine and aggressive male reproductive partners who would be more sociable and inclined to provide paternal investment.

Regards this latter suggestion, longstanding investigation suggests women’s mating and partnering decisions often involve adaptive trade-offs between masculine ‘good genes’, and high paternal investment from less-masculine male partners (Gangestad & Simpson, 2000; Kruger, 2006; Little, Connely, Feinberg, Jones, & Roberts, 2011; Quist et al., 2012; Trivers, 1972). Preference for robust masculinity is thought to be enhanced under high pathogen presence (DeBruine, Jones, Crawford, Welling, & Little, 2010; Little, DeBruine, & Jones, 2010), elevated social inequality (Brooks et al., 2010), and constrained food resource availability (Gleeson & Kushnick, 2018). Consistent with findings that women prefer more masculine men for short-term liaisons (Little, Cohen, Jones, & Belsky, 2007; Little et al., 2011), women thought to be less reliant upon paternal investment also prefer more masculine male faces (Marcinkowska et al., 2019). Other research has shown that women with prior exposure to male violence, whether directed towards themselves (Borras-Guevara, Batres, & Perrett, 2017a, 2017b), or at other women (Li et al., 2014), prefer relatively less-masculine partners. Taken together, the above moderators of female preference suggest a range of context-specific adaptive influences that will moderate selection for, or against, masculinity, wherever women can exercise effective sexual autonomy.

Work by other authors can be seen to support selective modes similar to the ‘social benefits’ (Cieri et al., 2014; Hare, 2017; Sánchez Villagra & van Schaik, 2019) and ‘execution’ (Wrangham, 2014, 2018, 2019a, 2019b) hypotheses of human self-domestication. For example, multiple contributions consider the adaptiveness of enhanced human sociability and collaborative capacities, especially via cooperative reproductive effort (Burkart et al., 2014; Gettler, 2010; Hawkes, 2013; Hrdy, 2009; Sterelny, 2011), but also through food sharing at times of need (Dyble et al., 2016; Gurven, 2004; Gurven & Hill, 2009). With these perspectives in mind, we would expect more-sociable group members to elicit higher levels of support, thus receiving fitness advantages over less-capable co-operators. Alternatively, social mechanisms involving collective ostracism and capital punishment of excessively dominating or antisocial individuals would also limit the reproductive success of aggressive predispositions (Boehm, 2012, 2014; Wrangham, 2018, 2019a, 2019b). Further, beyond these intragroup selective mechanisms, cooperation in *inter*-group hostility is also thought to have promoted human sociability by eliminating lineages with lower collaborative ability (Alexander, 1990; Flinn et al., 2005; Kissel & Kim, 2018; Wrangham & Glowacki, 2012).

In a recent article, Richard Wrangham (2019a) reviews nine possible modes of evolutionary selection against reactive aggression in humans; including processes not previously associated with human self-domestication. Each of these might act to promote greater sociability via reduced physiological masculinity, either directly (by suppressing reactive aggression), or indirectly (by favouring collaborative sociability). As in previous contributions to the topic, Wrangham (2014, 2018, 2019b) argues for the primacy of group collaboration (specifically, ‘language-based conspiracy’) in the execution of domineering and aggressive ‘alpha males’—presumably the most masculine male in the local population. Whilst helpfully extending current literature, this contribution may also serve to highlight some limitations needing further theoretical development. Given its focus, I discuss this work in some detail throughout the remainder of this section.

Sánchez Villagra and van Schaik (2019) recently argued that lack of a specific ‘time window’ is a hurdle to be overcome if the self-domestication hypothesis is to be truly testable. In effect, any continuing discourse requires a categorical clarification regards the purported timeline (or, *timelines*) during which self-domestication is thought to have occurred. However, whilst empirical and theoretical studies should clarify the period being considered, there is no a priori reason to assume self-domestication took place at only a single point in human evolution. In domesticated animals, diagnoses of domestication syndrome are made in comparison to a non-, or relatively less-, domesticated reference. Relevant symptoms should be apparent wherever there is continuing selection against aggressive reactivity. Thus, to satisfy the requirements of scientific testability, empirical studies must specify the period of interest, and the comparative samples to be examined. Where self-domestication is hypothesised, we should expect evidence of a syndrome of behavioural, physiological, and morphological changes across a given a period of evolutionary development, from one comparator to the other.

Hare et al. (2012) have convincingly documented the effects of self-domestication among wild bonobos, whose socioecological context allows coalitions of females to thwart non-consensual sexual advances—even from physically dominant males. Clark and Henneberg (2015, 2017) diagnosed similar ‘wild’ self-domestication as a factor in the evolution of *Ardipithecus ramidus*, around the dawn of the hominin lineage. Cieri et al. (2014) focussed their analysis on craniofacial feminization in Homo sapiens over the past 200,000 years. Elsewhere, Hare (2017) has inferred effects from self-domestication across three periods of human evolution: (1) the emergence of *Homo erectus*; (2) as a driver of increased brain size over the past 2 million years; and (3) as an explanation for the appearance of ‘behavioural modernity’ from around 50 thousand years ago. Such a diversity of instances implies a range of factors that might act against reactive aggression, to greater or lesser extent, at various periods throughout human pre-history—as well as in non-human vertebrate taxa. Wrangham’s (2019a) latest analysis focusses on selection against reactive aggression in a pre-sapiens ancestor—he suggests, most likely *Homo heidelbergensis*—and frames self-domestication as the reason for the emergence of *Homo sapiens* from this archaic lineage, around 300,000 years ago. He describes a vision of language-using sub-alpha male conspirators acting in concert to prevent reproductive monopolisation of women by any aggressive alpha male. Oddly, however, this depiction seems to imply pre-existing capacities for exactly the type of sociable cooperation (Cieri et al., 2014; Hare, 2017; Wrangham, 2019b) and language ability (Benítez-Burraco & Kempe, 2018; Thomas & Kirby, 2018) that self-domestication is often intended to explain. Given these attributed features, we might logically assume an earlier process of self-domestication in our lineage; however, ‘language-based’ conspiracy seems unlikely to have played a role much before this time.

Wrangham (2019a) argues against any effect from female masculinity preferences or mate-choice during the period in question, asserting that women would be unable to resist a determined alpha male, so could not sufficiently dampen the reproductive fitness of masculine aggression. However, fitness among aggressive males does not equate directly to the sum of forced copulations, or even pregnancies—this especially in humans, where male endocrine regulatory regimes point to longstanding strategies of paternal investment and care (Gettler, 2010; Gettler, McDade, Feranil, & Kuzawa, 2011). Boothroyd et al. (2017) recently showed that both low and high extremes of paternal masculinity in a traditional society are associated with reduced offspring survival (hence lower fitness) compared to moderately masculine men. One interpretation of these findings suggests they reflect theorised trade-offs between robust masculine genes and supportive paternal investment—implying fitness is optimised at the midpoint of any human masculinity continuum. Thus, whilst pre-historic ‘alpha’ males might maximise forced copulations, this strategy should not be assumed to translate to elevated fitness. Established reproductive advantages from shared care of infants favoured the evolution of alloparenting among related women, but must also have promoted paternal and alloparental care in men (Hrdy, 2009). Alloparenting is thought to have enhanced the sociability adults and promoted care-eliciting paedomorphism in infants (Hrdy, 2009); implying both selection in favour of sociable temperament, as well as juvenilised morphology reminiscent of domestication syndrome.

On a related point, whilst human female choice is conventionally examined as the preferences of individual women, this narrow formulation neglects any potential for cumulative social influence from such preferences. In this regard, evaluating ‘language-based conspiracy’ as a distinct alternative to ‘female choice’ subtly obscures the possibility of any conspiracy among human females; however, there is no necessary dichotomy between conspiracy and female choice as drivers of human self-domestication—unless one subscribes to a prior assumption that only male actors would be motivated to collaborate against domineering and violent men. Wrangham (2019a) may acknowledge this point when he states that ‘female choice *alone’* [my emphasis] is an unlikely driver of human self-domestication at this time.

Given domestication syndrome in humans presents as a relative ‘feminization’ of the phenotypic mean (Cieri et al., 2014; Wrangham, 2019b), and assuming self-domestication proceeded by selection against particularly-masculine aggressive reactivity, it seems logical to expect human females have long shown a heightened predisposition towards the behavioural typology of self-domestication—that is, *relatively* lower aggressive reactivity and an enhanced capacity for sociable and collaborative interaction. With this in mind, we might consider an illustrative alternative to the multi-male group of sub-alpha conspirators, one characterised by capable female co-operators—possibly with a post-reproductive grandmother or two—and a succession of socially-intolerant reactively-aggressive ‘alpha’ males. In this situation, a dominant male would need to be physically masculine enough to defeat other aggressive male challengers, but the most likely perpetrators of any conspiracy (language-based or not) would be women; relatively more-sociable women; with or without a familial bond; but, undoubtedly, with a shared reproductive interest in the promotion of alloparenting and paternal support for their progeny. The point of this illustration is not to suggest that female choice alone has promoted human self-domestication, but to highlight that both male and female reproductive strategies must be considered with regard to selection for or against masculine aggression in human evolution.

In light of evidence among contemporary humans, Wrangham’s (2019a) decidedly male language-based conspiracy must have been an influence in recent human self-domestication; but it could not have emerged until substantial sociability and language capacity had already appeared in men. As such, it seems unlikely to have been the only factor acting in favour of human self-domestication. Other influences must already have been in operation, or else, given the sociability and language capacities inferred, would likely have emerged around the same period. As discussed in Gleeson and Kushnick (2018), any pre-historic selective mechanism proposed to have promoted sociability and self-domestication in humans must always have operated in tandem with complimentary processes and in opposition to any contrasting effects. It follows that contributing drivers need not be approached as competing hypotheses; nor should we expect to identify a singular overall cause of human self-domestication. Certain factors may be more prominent at particular periods, but relative influence would seem difficult to test or quantify with scientific precision. We can assume some factors, such as language-based conspiracy, would emerge only after human sociability and communicative capacity were already sufficiently enhanced by prior group interaction and selection for elevated social tolerance.

Discussing noted decline in human sexual dimorphism since the mid-Pleistocene, Frayer and Wolpoff (1985, p. 462) suggest ‘simplistic, single cause models’ are unlikely to provide an explanation. Similarly, Sterelny (2012, p. 75) eschews ‘key-innovation’ models of human sociability, instead citing multiple factors likely to act as positive feedback for cooperative ability in humans. If we assume self-selection against aggression was a cumulative result of daily socio-sexual interaction over hundreds, if not thousands, of millennia across a variety of evolving hominin taxa, this seems a logical expectation. In this vein, Table 3 summarises several known or theorised factors likely to have influenced human self-domestication, presenting them as positive and negative feedback to emphasise interactions and tensions between them. Each is likely to have varied over deep-time in response to differing socioecological contexts, and stages of human physical, cognitive, and social evolution.

**Table 3.** Selective feedbacks affecting physiological masculinity, sociability, and self-domestication

| Feedback | Relevant influences |
| --- | --- |
| **Positive** | Increased altriciality and alloparenting<sup>1</sup> |
|  | Female preference for paternal investment<sup>2,3</sup> |
|  | Increased reliance on social learning and a cooperative sociocultural niche<sup>4-8</sup> |
|  | Group ostracism or execution of aggressive individuals<sup>9-12</sup> |
|  | Group collaboration in inter-group hostilities<sup>13,14</sup> |
| **Negative** | Direct and physical male-male contest competition<sup>15,16</sup> |
|  | Female selection in favour of masculine ‘good genes’<sup>17,18</sup> |

Notes: <sup>1</sup>(Hrady, 2009); <sup>2</sup>(Cieri et al., 2014); <sup>3</sup>(Kruger, 2006); <sup>4</sup>(Cieri et al., 2014; Ellis, 2016; Hare, 2017; Sterelny, 2011, 2018); <sup>5</sup>(Ellis, 2016); <sup>6</sup>(Hare, 2017); <sup>7</sup>(Sterelny, 2011); <sup>8</sup>(Sterelny, 2018); <sup>9</sup>(Boehm, 2014); <sup>10</sup>(Wrangham, 2018); <sup>11</sup>(Wrangham, 2019a); <sup>12</sup>(Wrangham, 2019b); <sup>13</sup>(Alexander, 1990); <sup>14</sup>(Wrangham & Glowacki, 2012); <sup>15</sup>(Carrier & Morgan, 2015); <sup>16</sup>(Hill et al., 2017); <sup>17</sup>(Holzleitner & Perrett, 2017); <sup>18</sup>(Marcinkowska et al., 2019).

## 8. Outstanding questions and further research

Associated behavioural, physiological, and morphological changes under domestication have been repeatedly observed—and experimentally demonstrated—in multiple vertebrate populations over a significant period (Agnvall et al., 2017; Belyaev, 1979; Darwin, 1868; Hemmer, 1990; King & Donaldson, 1929; Kruska, 1996; Leach, 2003; Trut, 1999; Trut et al., 2009). Processes maintaining these associations, and their operation under natural and cultivated conditions, are the subjects of recent and continuing research effort (Hare, 2017; Hare et al., 2012; Sánchez-Villagra et al., 2016; Theofanopoulou et al., 2017; Wilkins, 2017, 2019; Wilkins et al., 2014; Wrangham, 2019a). To expand physiological understanding, further study of mechanisms effecting the hypoplasia of NCC-derived tissues among animal domesticates should prove particularly useful.

On the expectation that most features of domestication syndrome derive from dampened NCC migration, research on piebaldism seems a particularly worthwhile area of focus. The lack of widespread piebald depigmentation in humans (other than as a noted and rare pathology) presents an open question for human self-domestication research. To my knowledge, there is no suggestion that human piebaldism results from domestication-related mechanisms. It may be useful to investigate whether genetic markers of piebaldism are more common in humans today than ancestrally, or to examine different frequencies of occurrence between contemporary human populations and cultures. Note that gradated variation in skin melanin content between recent human populations is unrelated to any predicted effect of self-domestication since these reflect melanosome and melanin variability, rather than differences in the distribution of NCC-derived melanocytes (Lin & Fisher, 2007).

A range of recent sources consider the genetics of domestication, providing solid foundations for further investigation (e.g. Benítez-Burraco, Lattanzi, & Murphy, 2016; Benítez-Burraco, Pietro, Barba, & Lattanzi, 2017; Benítez-Burraco, Theofanopoulou, & Boeckx, 2016; Fallahshahroudi et al., 2018; Løtvedt, Fallahshahroudi, Bektic, Altimiras, & Jensen, 2017; Theofanopoulou et al., 2017; Wilkins, 2017; Wilkins et al., 2014; Wright, 2015). Comparative studies have identified multiple genes repeatedly altered in domesticated lineages, several of which are thought to effect the function of NCCs (Theofanopoulou et al., 2017; Wilkins, 2017). Links between NCCs and masculine features, surveyed here, suggest future studies could benefit from a focus on sexual differentiation and masculine development, including any influence from androgen receptors and KIT ligand functioning on embryonic NCC migration. Previous research has proposed relationships between human self-domestication and various pathologies associated with brain language centres, suggesting autism spectrum disorder and schizophrenia might both be affected by domestication-related processes (Benítez-Burraco, 2017; Benítez-Burraco, Lattanzi, et al., 2016; Benítez-Burraco et al., 2017). Craniofacial hypermasculinisation associated with autism spectrum disorder (Tan et al., 2017)—a syndrome disproportionately present in human males—could prove relevant in light of links between NCCs and masculine phenotypes described here.

Present constraints mean that the examination of NCC-derived masculine traits presented in Section 3 is far from exhaustive; a more extensive and systematic comparison might prove productive. NCC pluripotency and the diversity of lineage-specific features derived from them, testifies to their elevated mutagenic capacity. The regularity with which these cells act as progenitors for masculine features is likely due to the frequency of evolutionary differentiation based on male sexual signalling and competition (Lande, 1981; McCullough, Miller, & Emlen, 2016; Tinghitella et al., 2018; Turner & Burrows, 1995), rather than any conserved predisposition to influence masculine physiology per se. Having said this, any influence from androgen receptors upon NCC migration would suggest the latter is a possibility. General associations between NCCs and vertebrate masculinity might theoretically link masculine signals to immune system functions via NCC-derived features such as the thymus and immune components of bone marrow (Foster et al., 2008; Han et al., 2015; Komada et al., 2012). Such a link might inform further research into sexual differences in immune function and pathology, possibly extending the handicap principle (Zahavi, 1975; Zahavi & Zahavi, 1999) and the ‘testosterone as immune suppressant’ hypothesis (Folstad & Karter, 1992). This being the case, an increased focus on the roles, and interactions, of NCCs and androgen receptors—as opposed to simple testosterone levels—in male signalling and immune function would likely prove insightful.

The present article has examined documented pre-historic reduction in human masculinity, with the expectation—after Cieri et al. (2014)—that decline in masculine physiology and aggressive behavioural predispositions played important roles in enhanced human sociability, language capacity, and sociocultural sophistication. However, this focus on masculinity should not preclude related and important changes occurring among hominin females. Domestication syndrome is known to reliably influence specifically female physiology, suggesting areas for further productive investigation; perhaps especially with regard to heterochronic change known to affect seasonality and reproductive timing, with potential implications for lifecycle fertility and human menopause.

## 9. Conclusion

Repeated developmental links between embryonic NCCs and a wide range of vertebrate male secondary sexual features imply selection against masculine-typical traits (including elevated reactive aggression) will influence NCC-migration more generally, thereby promoting domestication syndrome in a given lineage. In light of NCC contributions to masculine features in humans, pre-historic reduction in human sexual dimorphism is consistent with a process of human self-domestication. Regarding the NCC (Wilkins et al., 2014) and androgen (Cieri et al., 2014) hypotheses of domestication, and human self-domestication, repeated associations between NCCs and vertebrate masculine traits implies cellular hypoplasia due to dampened NCC migration would be sufficient to cause domesticated reductions in masculinity and sexual dimorphism. However, androgen receptors conspicuously moderate the growth and development of masculine features and may influence NCC migration— though this latter effect requires further investigation.

Self-domestication has been characterised as a necessary process in human evolution towards a more cooperative and sociable form; allowing the development of human culture, language, and complex knowledge sharing. Expansion of cultural complexity required a substantially enhanced capacity for social interaction and collaboration. Although the mechanisms proposed to explain these features vary between interested authors, any substantial increase in sociability must logically imply some fitness benefit and a level of selection in its favour. A range of variable and interacting influences likely effected this process throughout different periods of our evolutionary development. Enhanced cooperative capacity has facilitated human adaptation to the uniquely complex sociocultural niche we occupy today. Further self-domestication research will extend current understanding of its effects on human biology and behaviour, as individuals, and as a global society.

## Acknowledgements

I would like to acknowledge a debt of gratitude to Geoff Kushnick, Colin Groves, Katherine Balolia, and Heloisa Mariath for their welcome advice, encouragement, and support during the production of this manuscript. This work was supported by an Australian Government RTP scholarship.

